# Effects of Experimental Design, Genetic Architecture and Threshold on Power and False Positive Rate of Genome-Wide Association Studies

**DOI:** 10.1101/2022.02.19.481168

**Authors:** Zhi Loh, Sam Clark, Julius H. J. van der Werf

**Author notes:** Corresponding Author: Zhi Loh, Flat 14 Room 2, Wright College and Village, University of New England, Armidale NSW 2351 Australia. These authors contributed equally to this work.

## Abstract

Genome-Wide Association Studies are an important tool for identifying genetic markers associated with a trait, but it has been plagued by the multiple testing problem, which necessitates a multiple testing correction method. While many multiple testing methods have been suggested, e.g. Bonferroni and Benjamini-Hochberg’s False Discovery Rate, the quality of the adjusted threshold based on these methods is not as well investigated. The aim of this study was to evaluate the balance between power and false positive rate of a Genome-Wide Association Studies experiment with Bonferroni and Benjamini-Hochberg’s False Discovery Rate multiple testing correction methods and to test the effects of various experimental design and genetic architecture parameters on this balance. Our results suggest that when the markers are independent the threshold from Benjamini-Hochberg’s False Discovery Rate provides a better balance between power and false positive rate in an experiment. However, with correlations between markers the threshold of Benjamini-Hochberg’s False Discovery Rate becomes too lenient with an excessive number of false positives. Experimental design parameters such as sample size and number of markers used, as well as genetic architecture of a trait affect the balance between power and false positive rate. This experiment provided guidance in selecting an appropriate experimental design and multiple testing correction method when conducting an experiment.

## Introduction

Since high-density genotyping arrays using abundant genetic markers such as Single Nucleotide Polymorphisms (SNPs) have become available, Genome-Wide Association Studies (GWAS) has become an important tool in gene discovery (Wang and Xu 2019). Hundreds of thousands to several millions of genetic markers can now be used in association studies, where the aim is to estimate and test the effect of genetic variants linked to a Quantitative Trait Locus (QTL). This has provided a huge research opportunity but the use of large numbers of markers to be tested has also introduced a multiple testing problem of an unprecedented scale. Multiple testing significantly increases the number of false positives when using a standard significance threshold, thus necessitating the use of a correction method to adjust this threshold (Tam *et al*. 2019).

The most popular method of controlling the number of false positives is the Bonferroni correction. This multiple testing correction method is based on the joint distribution of all the Student’s t-distribution for each individual linear contrast, with the assumption that each of these tests are independent to one another (Dunn 1961). This method had gained popularity due to its simplicity (Ionita-Laza *et al*. 2013; Llinares-López *et al*. 2015), and is considered one of the most effective methods in controlling the number of false positives (Wilson 2019). However, the Bonferoni method has also been criticized when applied to GWAS as with very large numbers of SNPs tested, it has been perceived as being overconservative, leading to reduced power in identifying causal variants (Wilson 2019; Gao *et al*. 2010; Huang *et al*. 2018; Llinares-López *et al*. 2015). The situation has been further exacerbated by decreasing cost of genotyping, and it now has become common practice to use all genetic variants obtained from Whole Genome Sequence (WGS) information, often exceeding 25 million marker genotypes per sampled individual (Huang *et al*. 2018; Tam *et al*. 2019; Visscher *et al*. 2017).

Alternative multiple testing correction methods have been introduced, many of which have reduced stringency. One class of alternatives is those methods that attempt to control the False Discovery Rate (FDR), with one of the most popular methods being the Benjamini-Hochberg’s False Discovery Rate (BH-FDR) method. Initially introduced by Simes (1986), this method aims at testing the ranked p-values against a stepwise threshold that varies based on the rank of the p-values, with the most significant p-value subjected to the most stringent threshold, and other p-values that are less significant are subjected to more lenient threshold. An example of its implementation is provided in Table 1.

**Table 1.** An example of implementation of BH-FDR. For this example, 10 SNPs have been tested and have their p-values calculated according to their rank j. The point where the p-value of the marker falls below of that calculated from the stepwise threshold is at *j* = 4 is, and this is the point where the threshold of BH-FDR is set. Note that only the most significant marker (i.e. *j* = 1) had been subjected to the stepwise threshold equivalent to a Bonferroni correction.

| Index of ranked p-values ($j$) | Ranked p-values | Stepwise Threshold ($\frac{0.05 j}{n_{snp}}$) | Decision<br>(Accept or Reject Null) |
| --- | --- | --- | --- |
| 10 | 0.676 | 0.050 | Accept |
| 9 | 0.324 | 0.045 | Accept |
| 8 | 0.213 | 0.040 | Accept |
| 7 | 0.119 | 0.035 | Accept |
| 6 | 0.087 | 0.030 | Accept |
| 5 | 0.034 | 0.025 | Accept |
| 4 | 0.012 | 0.020 | Reject |
| 3 | 0.010 | 0.015 | Reject |
| 2 | 0.006 | 0.010 | Reject |
| 1 | 0.002 | 0.005 | Reject |

Many GWAS have chosen the BH-FDR multiple testing correction method on the grounds of overconservativeness of the Bonferroni correction, but appeared to have no consideration on the possibility of increased false positive rate. In the context of gene expression analysis, Huang *et al*. (2018) considered BH-FDR to have a better balance between power and false positive rate, although they also commented that the use of BH-FDR resulted in an inflated false positive rate whereas Bonferroni correction had a significantly lower number of false positives. Another consideration in most GWAS is that with the use of dense markers, marker genotypes can be highly correlated. Benjamini and Yekutieli (2001) suggested that in theory this method is valid even when the assumption of independence between tests is violated, as would be the case in GWAS based on dense marker genotypes An actual study on the ability of BH-FDR in controlling the false positive rate in a GWAS and the need to account for the lack of independence between tests is lacking however.

Several factors could impact the success in detecting QTL associated with a trait while controlling the false positive rate, including parameters related to the genetic architecture of the traits, i.e. the size of the QTL effects, and experiment design, most notably the sample size. Spencer *et al*. (2009) and Visscher *et al*. (2017) argued a reduced power of GWAS with small sample size, while Forstmeier *et al*. (2017) argued an increased false positive rate with small sample size in any statistical test. The low power alongside with increased false positive rate could have contributed to the low replicability of a GWAS experiment where hits from the previous studies failed to be replicated in subsequent studies (Heller and Yekutieli 2014; Wang and Zhu 2019). Spencer *et al*. (2009) and Visscher *et al*. (2017) argued that increasing the sample size is the most effective way to increase the power of GWAS, and while the number of positives increases with sample size, it is unclear how much of the positives are true positives (a summary of number of positives reported in previous publication is provided in Table 2).

**Table 2.** Summary of threshold used, sample size, and number of markers and positives used in previous publications. For studies that included multiple traits, data from only one trait was included. BH-FDR stands for Benjamini-Hochberg False Discovery Rate method, and BON for Bonferroni correction. The publications are ranked based on the sample size used.

| Publications | Sample Size | Number of Markers | Correction Method | Alpha | $-\log_{10}(\text{threshold})$ | Number of Positives |
| --- | --- | --- | --- | --- | --- | --- |
| Ghasemi *et al.* (2019) | 130 | 41,323 | BON | 0.05 | 5.91 | 7 |
| Zhang *et al.* (2013) | 319 | 48,198 | BON | 0.05 | 5.98 | 10 |
| Vanvanhossou *et al.* (2020) | 449 | 32,518 | BON | 0.05 | 5.81<sup>a</sup> | 4 |
| Signer-Hasler *et al.* (2012) | 1077 | 38,124 | BON | 0.05 | 5.88 | 8 |
| Xia *et al.* (2017) | 1141 | 677,855 | BON | 1.0 | 5.83 | 11 |
| Chang *et al.* (2018) | 1217 | 671,990 | BON | 0.05 | 7.12 | 11 |
| Al-Mamun *et al.* (2015) | 1449 | 48,640 | BON | 0.01 | 6.69 | 39 |
| Weerasinghe *et al.* (2019) | 3454 | 37974 | BON | 0.05 | 5.88 | 13 |
| Cai *et al.* (2019) | 5373 | 16,503,508 | BON | 0.05 | 8.52<sup>a</sup> | 58535 |
| Dakhlan *et al.* (2017) | 6463 | 48,599 | BON | 0.01 | 6.69 | 17 |
| Yin and König (2019) | 13,827 | 54,613 | BON | 0.05 | 6.04 | 10 |
| Jiang *et al.* (2019) | 294079 | 57,067 | BON | 0.005<sup>a</sup> | 7.00 | 15215 |
| Smolucha *et al.* (2021) | 155 | 49,204 | BH-FDR | 0.05 | 5.99 | 1 |
| Wang *et al.* (2017) | 880 | 51,727 | BH-FDR | 0.01 | 4.00 | 5 |
| Steri *et al.* (2019) | 946 | 135,992 | BH-FDR | 0.08 | 5.84 | 5 |
| An *et al.* (2020) | 1,217 | 67,192 | BH-FDR | 0.01 | 6.17<sup>a</sup> | 45 |
| Ibeagha-Awemu *et al.* (2016) | 1,246 | 76,355 | BH-FDR | 0.1 | 4.16<sup>a</sup> | 53 |
| Pegolo *et al.* (2020) | 1,369 | 23,173 | BH-FDR | 0.05 | 4.30 | 24 |
| Akanno *et al.* (2018) | 5,324 | 42,536 | BH-FDR | 0.10 | 3.16<sup>a</sup> | 294 |

<sup>a</sup> Values back-calculated using available data (i.e. alpha, number of SNPs, number of positives)

The aim of this study is to test the effects of multiple testing correction methods on the power and false positive rate of a GWAS experiment, and subsequently evaluate the effects of experimental design parameters and genetic architecture of a trait on the suitability of the methods. We use simulation to evaluate the power and false positive rate with Bonferroni and BH-FDR correction methods under varying parameter values.

## Methods

### Experiment Procedure

The effects of the GWAS parameters and multiple testing correction methods were evaluated using simulated genotypes and phenotypes. This simulation is conducted using Python (version 3.7.3). To simulate a GWAS experiment with independent markers, data from a genotype array with M markers (henceforth denoted as ***X***) was generated for N individuals. The distribution of the allele frequency of the markers following a symmetrical Beta distribution (i.e. *Beta*(*a, a*)). Values used for the shape parameter a from the beta distribution are provided in Table 3. Some of the markers were nominated as QTL, and an effect size was assigned to each of the markers. The effect sizes of the QTL are distributed based on the following gamma distribution:

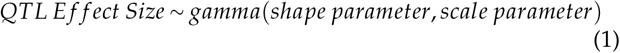

**Table 3.** Parameters tested in this study.

| Parameters | Default Values | Alternative Values |
| --- | --- | --- |
| Sample Size | 2000 | 200, 800, 1400, 3000, 5000 |
| Shape parameter for Distribution of Allele Frequencies ($\beta$) | 0.5 | 0.1, 0.2, 0.3, 0.7, 1.0 |
| Average QTL Effect Sizes ($\gamma$) | 0.4 | 0.1, 0.2, 0.3, 0.7, 1.0 |
| Number of Markers | 20k | 5k, 10k, 40k, 60k, 80k |
| Number of QTL | 100 | 20, 50, 300, 600, 1000 |

**Table 4.** The number of true positives (TP), correlated false positives (FPC) and uncorrelated false positives (FPU) under varying multiple testing correction methods and dependency between markers. Default parameters had been used in calculating the number of true and false positives for this table.

| Multiple Test-<br>ing Method | Bonferroni correction |  |  | BH-FDR |  |  |
| --- | --- | --- | --- | --- | --- | --- |
|  | TP | FPC | FPU | TP | FPC | FPU |
| Independent Markers | 7.53 | NA | 0.22 | 9.13 | NA | 0.76 |
| Correlated Markers | 7.86 | 51.12 | 0.80 | 12.23 | 98.44 | 10.18 |

The scale parameter is set at 1 for all simulation, while the shape parameter is varied based on the average QTL effect size, which would be provided in Table 3. The average QTL effect size is calculated as follow:

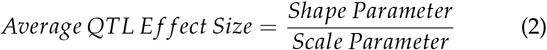

Markers that were not nominated as QTL would have their effect sizes marked at 0. Using the vector containing the effect sizes for all markers and QTL (denoted as ***a***), the additive genetic component of the phenotype (denoted as ***g***) is calculated as follows:

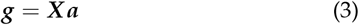

The residual component of the phenotype (denoted as ***e***) is then simulated using the variance of vector ***g*** and the narrow sense heritability of the trait *h*^2^. The residual component follows a normal distribution with mean of zero and variance as follow:

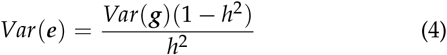

For all the parameter under study, the heritability was set at 0.3. The vector ***g*** and ***e*** were then summed to obtained the simulated phenotype of the individuals. A GWAS was then conducted using the genotype array and phenotype vector. Single SNP regression was used to estimate the effect sizes of the markers, which would then be used to calculate the p-values for each marker using the Student’s t-test.

Using the alpha = 0.05 for type 1 error, the thresholds from both Bonferroni correction and BH-FDR were calculated. The threshold for Bonferroni correction is defined as alpha divided by number of markers used in the experiment:

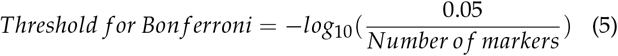

For BH-FDR in this experiment, the stepwise threshold is defined as follows (Simes 1986; Benjamini and Yekutieli 2001):

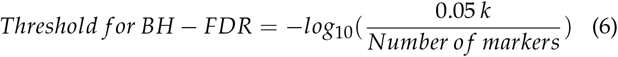

where *k* is the point where the *k*_*th*_ ranked -log(p-value) of the GWAS becomes larger than the stepwise threshold. The point is equivalent to the *j* = 4 from the example in Table 1. With these threshold, the number of true positives and false positives (henceforth denoted as #*TP* and #*FP* respectively) were recorded, from which the power and false positive rate, as well as differences between true and false positives (henceforth denoted as *ROC score*), were calculated. The power is defined as follows:

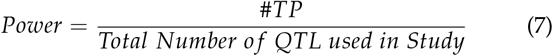

For the calculation of power only the QTL with effect size exceeding 0.1 *σ*_*e*_ were taken into account. The false positive rate is as follows:

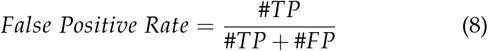

and the *ROC score* is defined as follow:

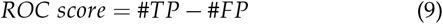

In this study the *ROC score* was used as a measure to test the capability of a threshold in balancing the power and false positive rate of a GWAS. This is equivalent to the weighted Youden ’s Index as described by Habibzadeh *et al*. (2016), who have utilized a Receiver Operating Characteristic (ROC) curve to establish the optimal threshold for clinical diagnostic tests.

A multiple testing correction method with its threshold having a high *ROC score* would be considered as capable of providing a better balance between power and false positive rates compared to another threshold with lower *ROC score*. A threshold with maximum *ROC score* would be considered as optimal. This is equivalent to having the point on the ROC curve where the tangent of the curve would be equal to 1, which has been demonstrated mathematically by Kaivanto (2008). The experiment was then repeated 200 times for each combination of parameter values.

To test the effect of correlations between marker genotypes on the optimal threshold and number of true and false positives, the experiment is repeated with pairwise marker linkage disequilibrium (denoted as *r*^2^) set at 0.8. This is achieved by copying the haplotype state of some of the alleles from one locus to its neighbouring locus while randomizing the haplotype state of other alleles, thus generating a genotype array with a controlled level of pairwise marker linkage disequilibrium. For correlated markers, besides the true positives, there were two types of false positives to be identified: (i) correlated false positives (*FPC*), defined as the false positives that its *r*^2^ more than 0.1 with one or more QTL, and (ii) uncorrelated false positives (denoted as *FPU*), defined as false positives that had its *r*^2^ below 0.1 with any of the QTLs. For the calculation of false positive rate, the number of FPU is used in place of #*FP* in equation (8) and (9).

### Parameters tested

A list of parameters and value tested is provided in Table 3. The default parameter values, alongside with their alternative values that were used in this study are provided in Table 3. When a parameter is under study, default values would be used of other parameters.

Besides the parameters listed in Table 3, the combined effects of sample size and number of markers on the power and false positive rate of GWAS were also tested. To test the combined effects of both parameters, additional simulations on variable sample sizes have been conducted with number of markers of 5k, 20k and 80k. The sample sizes used in this additional simulation are the same as those in Table 3 (i.e. N = 200, 800, 1400, 2000, 3000, 5000). This additional simulation is to test the power and false positive rate, as well as suitability of the multiple testing correction methods, of a GWAS experiment that involves small sample size but large number of markers, as in Steri *et al*. (2019) that have conducted a GWAS with 946 animals but with 135,992 markers.

## Results

### Parameters determining the threshold of multiple testing correction methods

#### Number of markers and sample size

The threshold from both multiple testing methods is influenced by the number of markers used in GWAS. With an increased number of markers used in GWAS, the threshold increases in stringency. This observation was made in both multiple testing correction methods in both independent and correlated marker system.

When Bonferroni correction is used, sample size does not have any effect on the threshold of GWAS. This is not the case for BH-FDR however, as the threshold from BH-FDR is significantly affected by sample size, with larger sample sizes decreasing the threshold. Generally, the threshold calculated by BH-FDR is less stringent than those calculated by Bonferroni correction (Figure 1).

**Figure 1.**
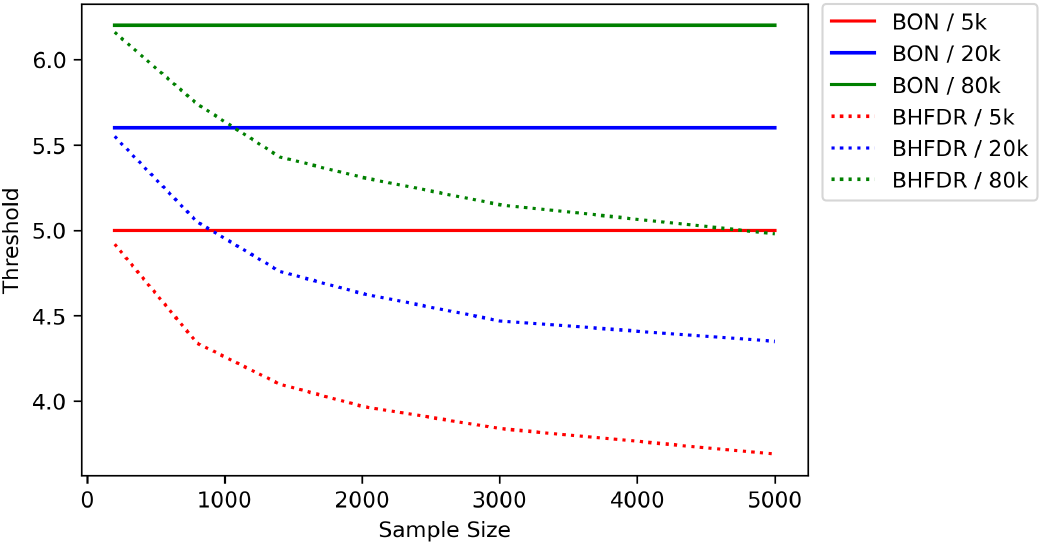
Threshold of Bonferroni correction (in solid lines) and BH-FDR (in dashed lines) under varying sample size and number of markers used in a GWAS experiment. This is the threshold under independent markers. The sample size is maintained at 2000, the number of QTL maintained at 100, and the average QTL effect sizes (*γ*) and allele frequencies (*β*) are at 0.4 and 0.5, respectively.

#### Number of QTL and QTL effects

The number of QTL does not have any influence on the threshold calculated from the Bonferroni correction. The number of QTL has an effect on the threshold of BH-FDR, however. When the number of QTL is small (e.g. 20) the threshold from BH-FDR approaches 4.9, and this threshold declines slightly to 4.63 with a number of QTL of 100, but then increases again gradually with larger numbers of QTL (Figure 2). A smaller average QTL effect sizes also increases the threshold slightly for BH-FDR (Figure 3). The allele frequency distribution does not have an effect on both multiple testing correction methods (Figure 4).

**Figure 2.**
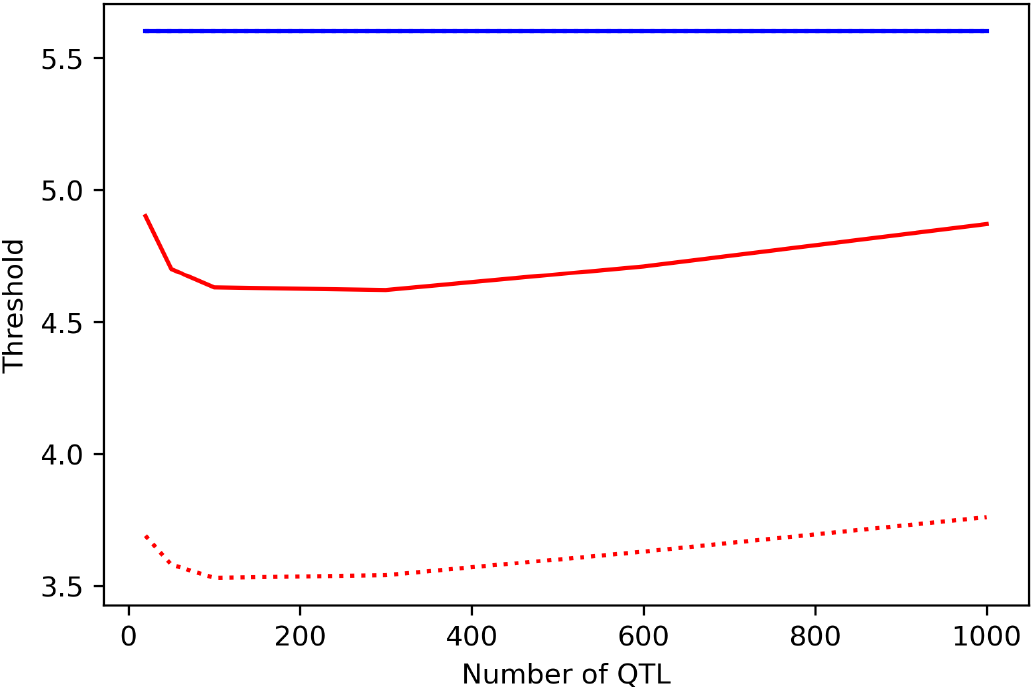
Threshold for Bonferroni correction (blue line) and BH-FDR (red lines) for both independent (solid lines) and correlated (dashed lines) markers with a varying number of QTL. The sample size was 2000, the number of markers 20k, the average QTL effect size (*γ*) set at 0.4 and shape parameter for allele frequencies distribution (*β*) set 0.5, respectively.

**Figure 3.**
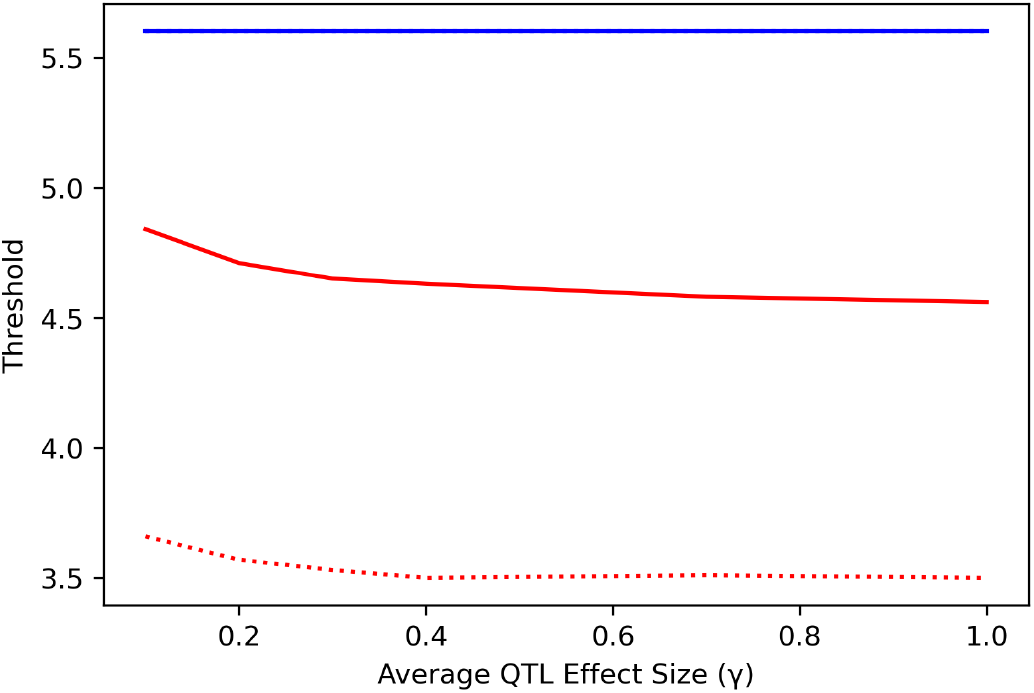
Threshold for Bonferroni correction (blue line) and BH-FDR (red lines) for both independent (solid lines) and correlated (dashed lines) markers under varying average QTL allele substitution effects (*γ*). The sample size used is 2000, with 20k markers and 100 QTL. The shape parameter of allele frequency distribution (*β*) is 0.5.

**Figure 4.**
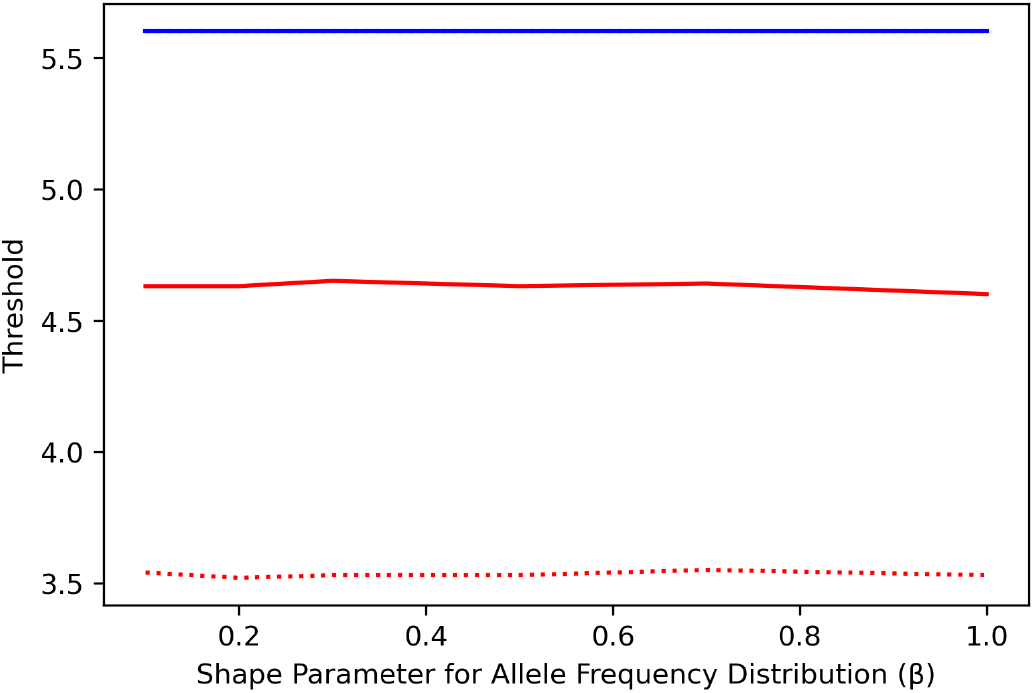
Threshold for Bonferroni correction (blue lines) and BH-FDR (red lines) for both independent (solid lines) and correlated (dashed lines) markers under varying shape parameter for allele frequency distribution (*β*). The sample size used is 2000, with 20k markers and 100 QTL, with average QTL effect size (*γ*) being set at 0.4.

#### Correlation between marker genotypes

For conventional Bonferroni correction, which assumes all markers are independent to one another, correlation between markers does not have any effect on the threshold for any of the parameters tested. For BH-FDR however, marker correlation has a significant effect on the threshold. Correlation between markers significantly decreases the GWAS threshold, which is evident from a reduced stringency in threshold for BH-FDR in Figure 4. With independent markers, the number of markers and sample size also have significant effects on the threshold of BH-FDR for correlated markers. While the trend is comparable with those in independent markers, the threshold calculated by BH-FDR is lower with correlated markers compared to independent markers for all marker numbers and sample sizes tested. Correlations between markers also caused a similar decline in BH-FDR threshold for all numbers of QTL, average QTL effect sizes and allele frequency distributions tested in this experiment.

### Parameters determining the power of GWAS

#### Number of markers and sample size

Due to an increased stringency in threshold from both multiple testing correction methods, the power decreases with an increased number of SNP markers used in a GWAS experiment. This observation was made for both independent markers and correlated markers. While correlations between markers increased the power for all marker number values tested, such an increase is more significant for experiments with a small number of markers, or when BH-FDR is used in GWAS (Figure 5).

**Figure 5.**
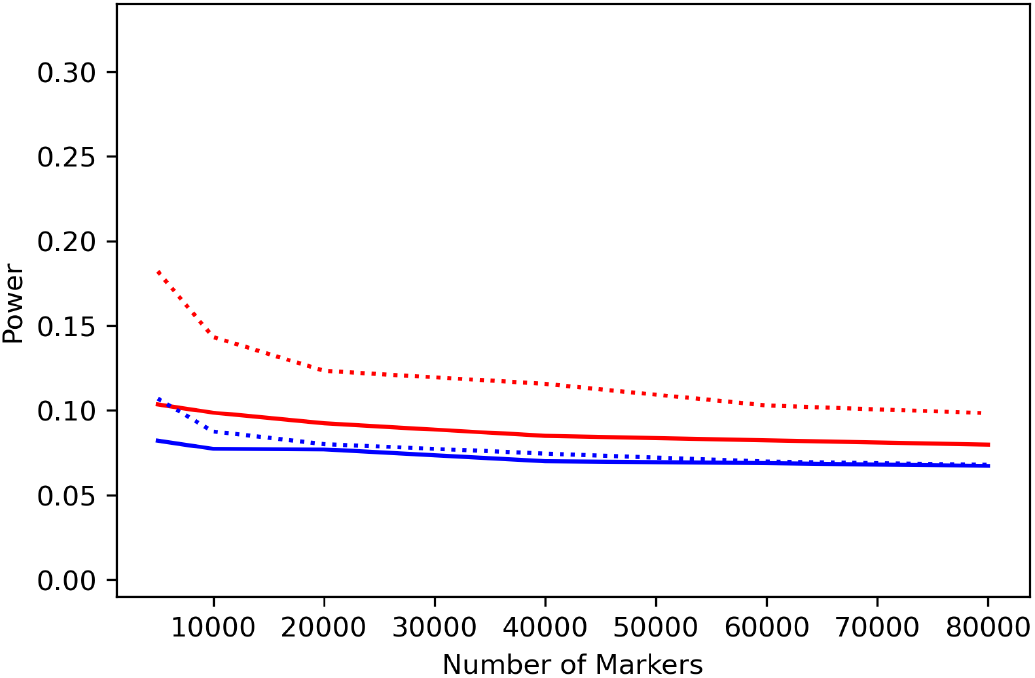
The power for Bonferroni correction (blue lines) and BH-FDR (red lines) under independent (solid lines) and correlated (dashed lines) markers under varying number of markers. The sample size is maintained at 2000, the number of QTL maintained at 100, the average QTL effect size (*γ*) set at 0.4 and shape parameter for allele frequencies distribution (*β*) set 0.5, respectively.

Increasing the sample size would increase the number of true positives and the power of GWAS, and this increase is more significant when BH-FDR is used. Correlation between markers has no effect on the power of GWAS if Bonferroni correction is used, but significantly increases the power for BH-FDR. This is attributable to an increased leniency in the threshold for BH-FDR with larger sample size (Figure 6).

**Figure 6.**
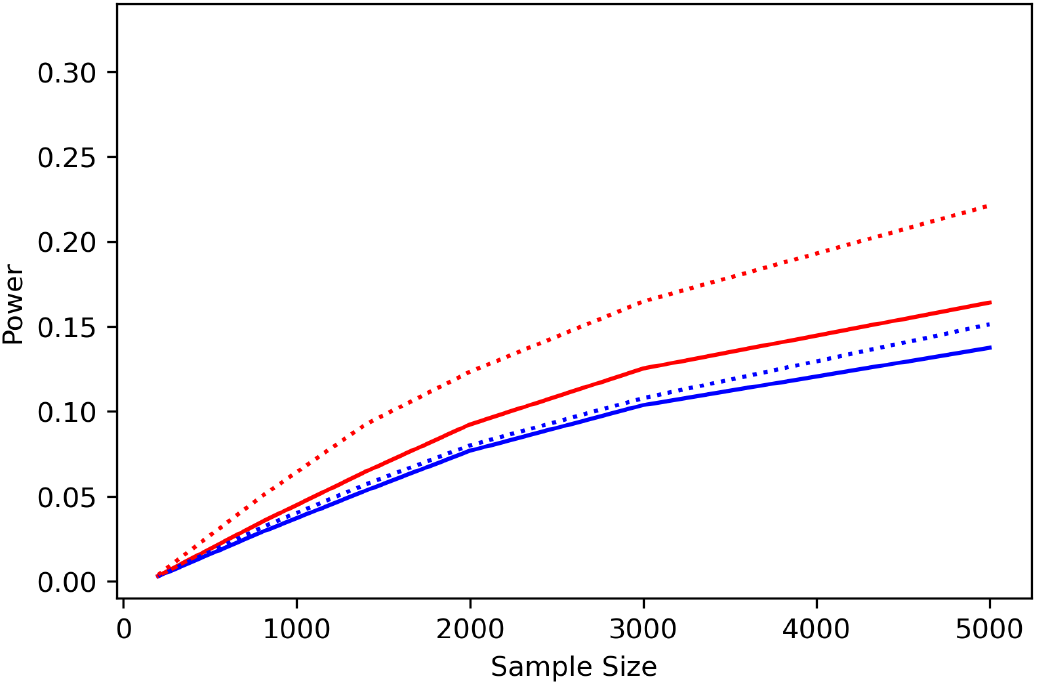
The power for Bonferroni correction (blue lines) and BH-FDR (red lines) for independent (solid lines) and correlated (dashed lines) for varying sample sizes. The number of markers and QTL are set at 20k and 100 respectively, the average QTL effect size (*γ*) maintained at 0.4 and shape parameter for allele frequencies distribution (*β*) set 0.5, respectively.

#### Number of QTL and QTL effects

The number of QTL that is associated with a trait has a significant effect on the power of detecting the QTL, with the power decreasing when the number of QTL increased, both for independent and correlated markers. This was observed for both multiple testing correction methods, although the power is higher for correlated markers when BH-FDR is used in the GWAS (Figure 7).

**Figure 7.**
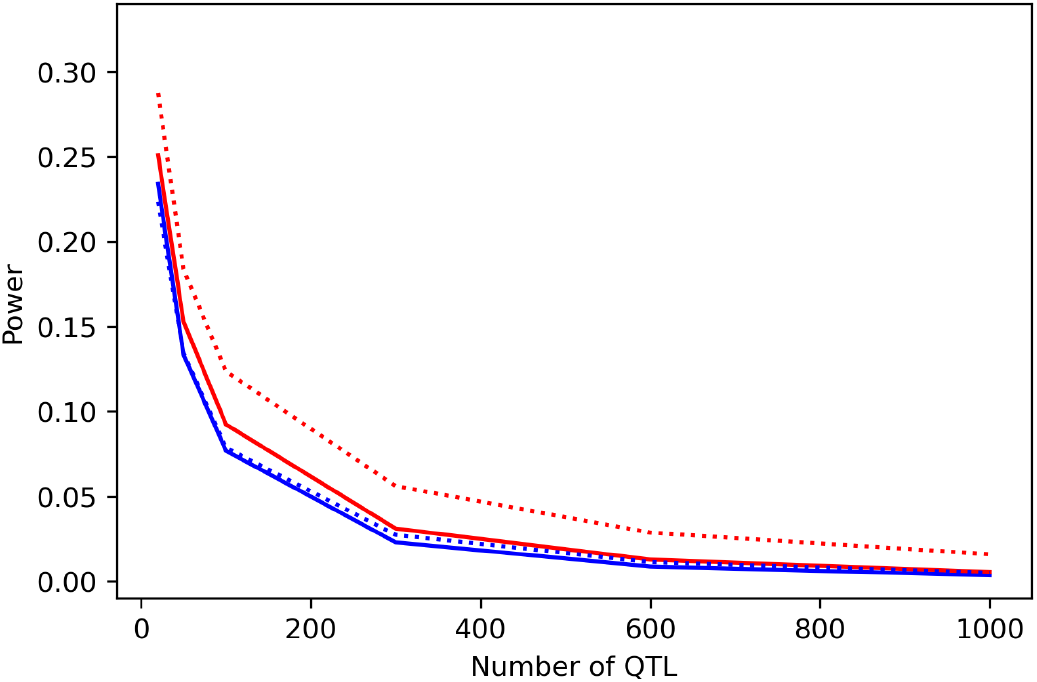
The power of Bonferroni correction (blue lines) and BH-FDR (red lines) for independent (solid lines) and correlated (dashed lines) under varying number of QTL. The sample size is maintained at 2000, the number of markers at 20k, with the average QTL effect size (*β*) set at 0.4 and shape parameter for allele frequencies distribution (*γ*) set 0.5, respectively.

The average QTL effect sizes (*γ*) has a significant effect on the number of true positives and power of GWAS. With an increased value of *γ*, the number of true positives increases until it starts to plateau by average QTL effect size of 0.4. In all cases, BH-FDR had a higher number of positives. Correlation between markers also increases the number of true positives for both multiple testing correction methods, and this increment is more significant for BH-FDR (Figure 8). The shape parameter for allele frequency distribution (*β*) has again no effect on the number of true positives and power of GWAS.

**Figure 8.**
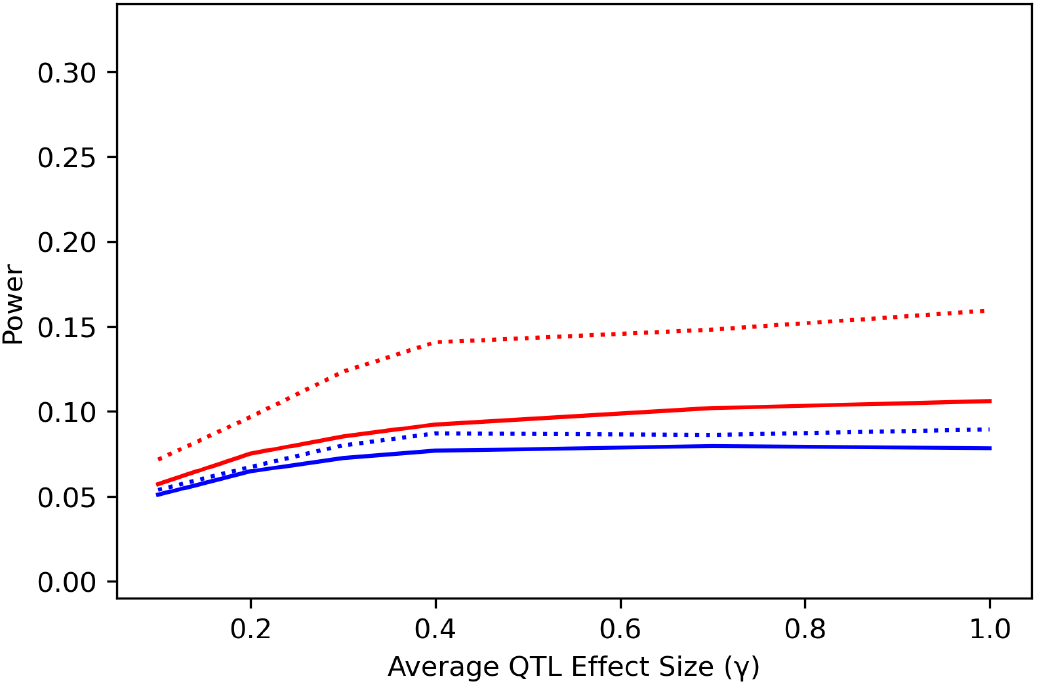
The power of GWAS under varying average allele substitution effects (*γ*). Blue lines represent results from Bonferroni correction and for red lines, BH-FDR. Solid lines represent independent markers and dashed lines represent correlated markers. The sample size is maintained at 2000, the number of markers and QTL at 20k and 100 respectively, and the shape parameter for allele frequency distribution (*β*) at 0.5.

### Effect of Parameters on False Positive Rate of GWAS

#### Number of markers and sample size

Despite the increasingly large number of tests needed to be conducted in a GWAS experiment with a larger number of markers, due to the increasingly stringent threshold, the raw number of false positives declines logarithmically. This is observed in both FPU and FPC. Due to a lower number of true positives associated with a more stringent threshold, however, the false positive rate is no longer significantly affected by the number of markers used in a GWAS experiment (Figure 9).

**Figure 9.**
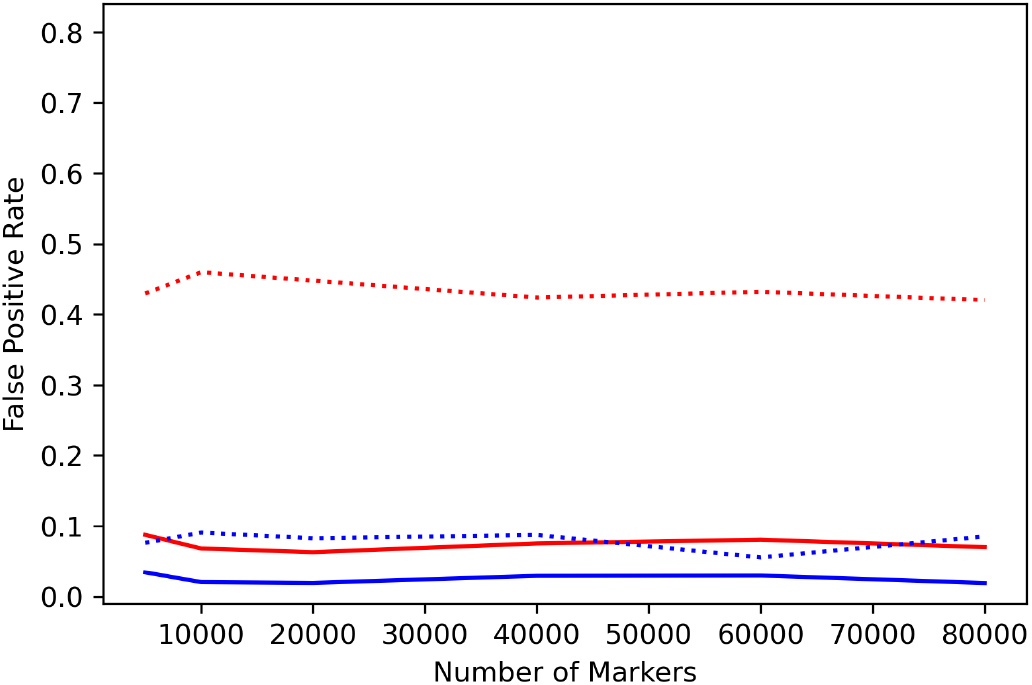
The false positive rate under varying number of markers used in a GWAS experiment. Blue lines represent Bonferroni correction and red lines represent BH-FDR. Solid lines represent independent markers and dashed lines represent correlated markers. The sample size is maintained at 2000, the number of QTL maintained at 100. The average QTL effect size (*γ*) is set at 0.4 and the shape parameter for allele frequency distribution (*β*) is set at 0.5.

Unlike marker number, sample size has a significant effect on the false positive rate of a GWAS experiment (Figure 10). The false positive rate increased significantly when the sample size is small (i.e. N=200). This trend was observed for both independent and correlated markers, and in both multiple testing correction methods. With larger sample size, the false positive rate remained relatively constant if the markers are independent. This is not the case for correlated markers however; the number of FPU increased significantly with larger sample sizes, and that led to an increase in false positive rate. While this was observed for both multiple testing correction methods, the false positive rate for BH-FDR is higher for all sample sizes tested in this experiment. While the number of markers does not have an effect on the false positive rate of a GWAS experiment under the default sample size (i.e. N=2000), for small sample size (N=200) a larger number of markers also strongly increased the false positive rate (Figure 11).

**Figure 10.**
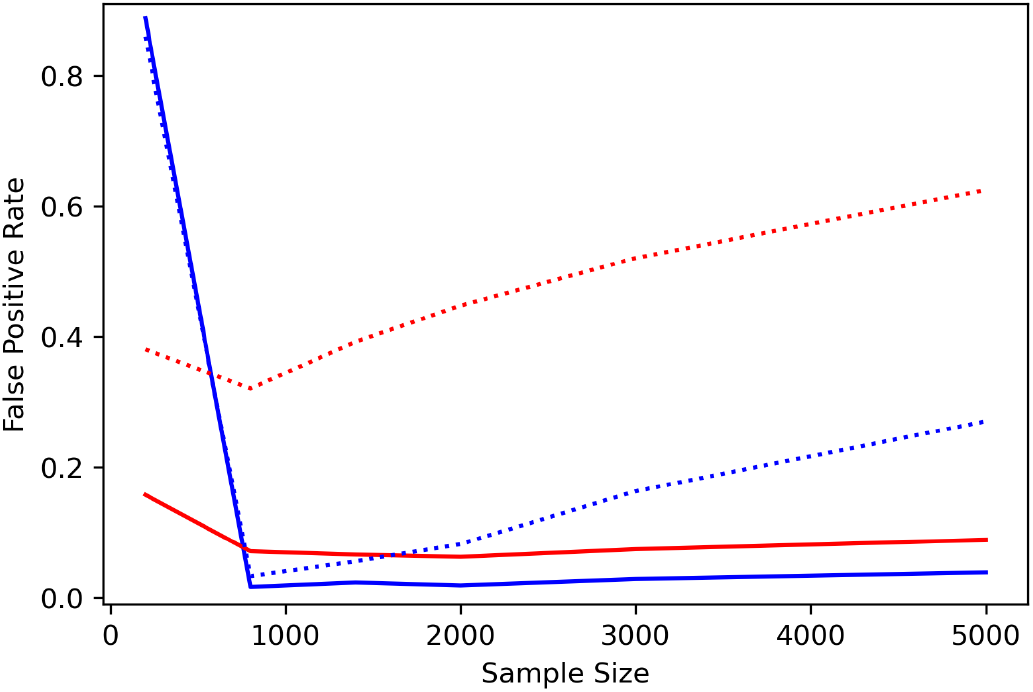
The false positive rate under varying sample size for Bonferroni correction (blue lines) and BH-FDR (red lines). Solid lines represent independent markers while dashed lines represent correlated markers. The number of markers and number of QTL are maintained at 20k and 100 respectively, and the average QTL effect size (*γ*) of 0.4. The shape parameter of allele frequency distribution (*β*) is set at 0.5.

**Figure 11.**
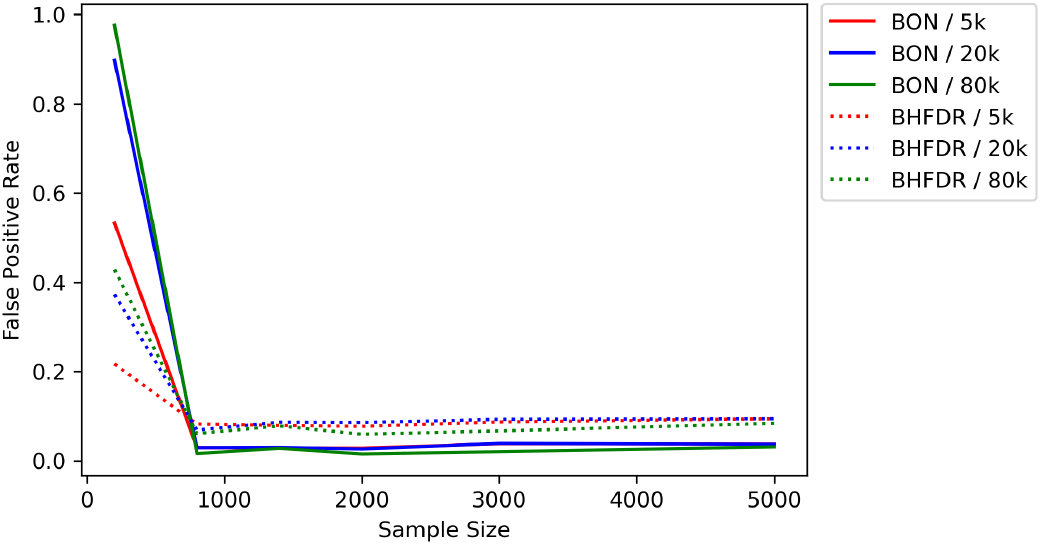
The effects of sample size on the false positive rate of GWAS under varying number of markers and correction methods. Solid lines represent the number of false positives for Bonferroni correction whereas dashed lines represent those of BH-FDR. The number of QTL is maintained at 100, and the average QTL effect sizes (*γ*) and allele frequencies (*β*) maintained at 0.4 and 0.5 respectively. Independent markers has been used in this plot.

#### Number of QTL and QTL effects

For independent markers, the false positive rate of a GWAS is not influenced by the number of QTL associated with a trait. This is not the case for correlated markers however; traits with small number of QTL with large effect sizes have a higher false positive rate compared to traits with large number of QTL with small effect sizes, and correlation between markers exacerbated that increment (Figure 12). This is caused by an increase in raw number of FPU and a decrease in raw number of true positives (as there is less QTL to be detected in the first place). The increase in the number of false positives with a small number of QTL is also due to an increase in significance from the increased proportion of variance explained by those QTL (which also explained the increase in power of GWAS with a small number of QTL). This increase in significance at the QTL also increases the significance of neighbouring null markers, thus increases the false positive rates.

**Figure 12.**
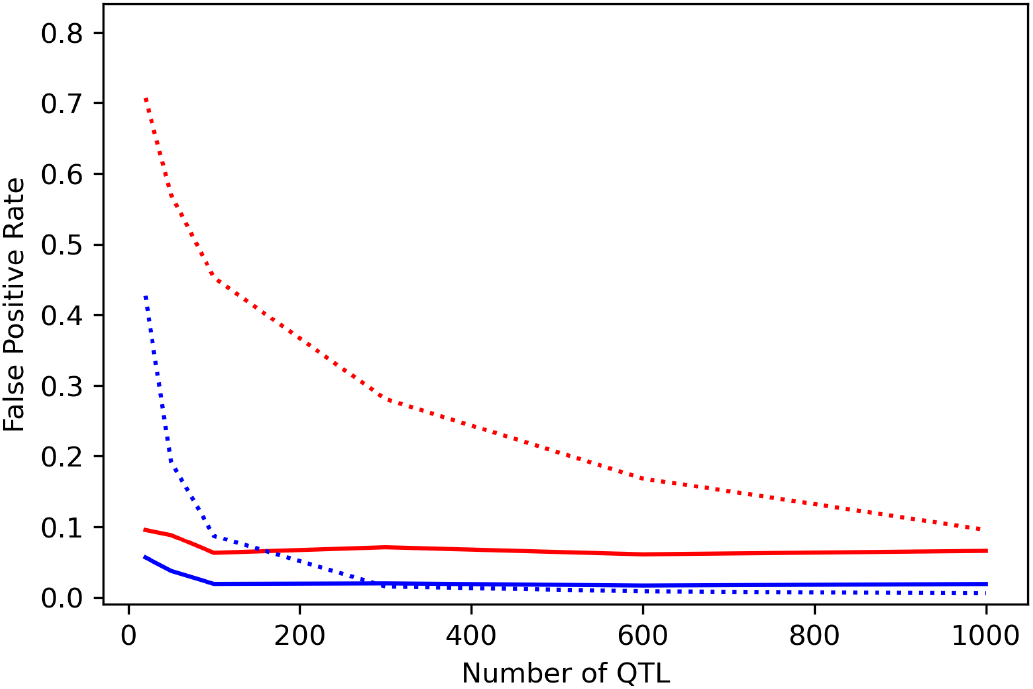
False positive rate associated with varying number of QTLs for Bonferroni correction (blue lines) and BH-FDR (red lines) and for independent (solid lines) and correlated (dashed lines) markers. The sample size is maintained at 2000, the number of markers at 20k, and the average QTL effect sizes (*γ*) and allele frequencies distribution shape parameter (*β*) maintained at 0.4 and 0.5 respectively.

The false positive rate is not significantly affected by the average QTL effect size (*γ*) when the markers are independent. But for correlated markers a lower value for *γ* significantly increased the number of FPU and false positive rate in both multiple testing correction methods. The number of false positives began to stabilize at an average QTL effect size of 0.4 (Figure 13). The number of false positives is not significantly affected by the distribution of the allele frequencies.

**Figure 13.**
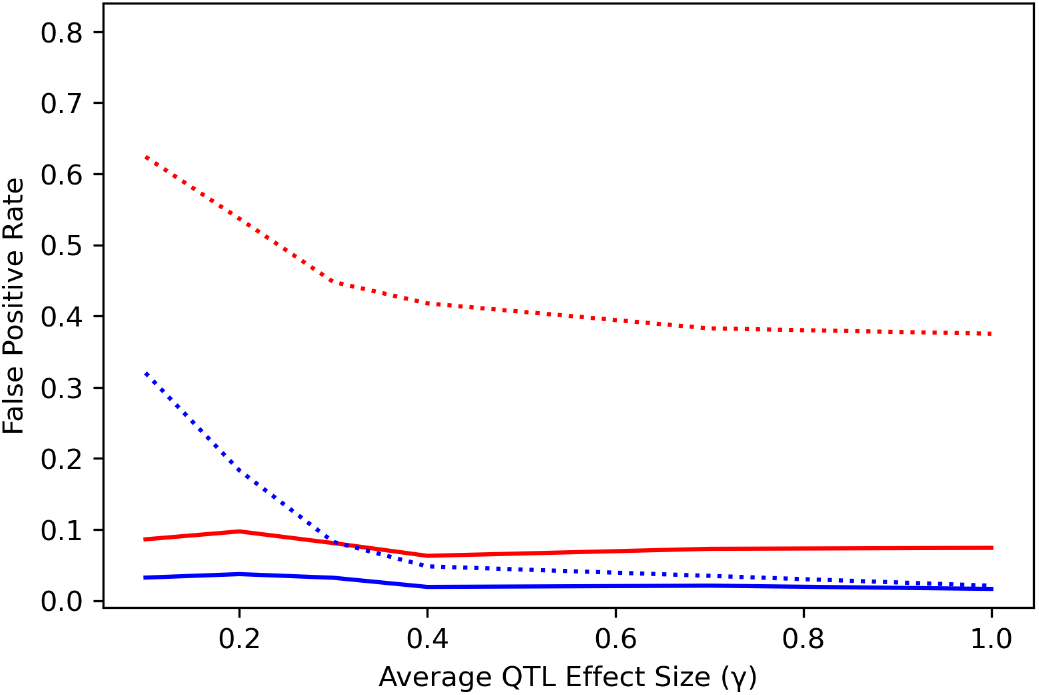
The false positive rate associated with varying average QTL effect size *γ* for Bonferroni correction (blue lines) and BH-FDR (red lines) and for independent (solid lines) and correlated (dashed lines) markers. The sample size is 2000, the number of markers and QTL 20k and 100, respectively, and the shape parameter of allele frequency (*β*) is 0.5.

#### Correlation between markers

For all the parameters tested, correlation between markers has a significant effect on the number of false positives detected in a GWAS. The presence of correlation significantly increased the number of false positives, although most of the false positives are correlated to the true QTLs (4). This observation can be made in both multiple testing correction method, although the numbers of both correlated and uncorrelated false positives are higher for BH-FDR compared to Bonferroni correction.

### Effects of Parameters on *ROC score* of Multiple Testing Correction Methods

With increasingly large numbers of markers used, there is a general decline in the difference between the True positive and the False positive marker (*ROC score*) for both multiple testing correction methods. For independent markers, BH-FDR had a higher *ROC score* compared to Bonferroni correction for all numbers of markers tested in this experiment. This suggests that the threshold for Bonferroni correction provided a less favourable balance between power and false positive rates in a GWAS experiment. The trend changes with the presence of correlations however; when the markers are correlated BH-FDR had a significantly reduced *ROC score* for all numbers of markers tested. This is attributable to an increased number of FPU when the assumption of independence is violated. With the exception of small number of markers used, which increases the *ROC score*, correlation between markers generally do not have a significant effect on the *ROC score* for the Bonferroni correction method (Figure 14).

**Figure 14.**
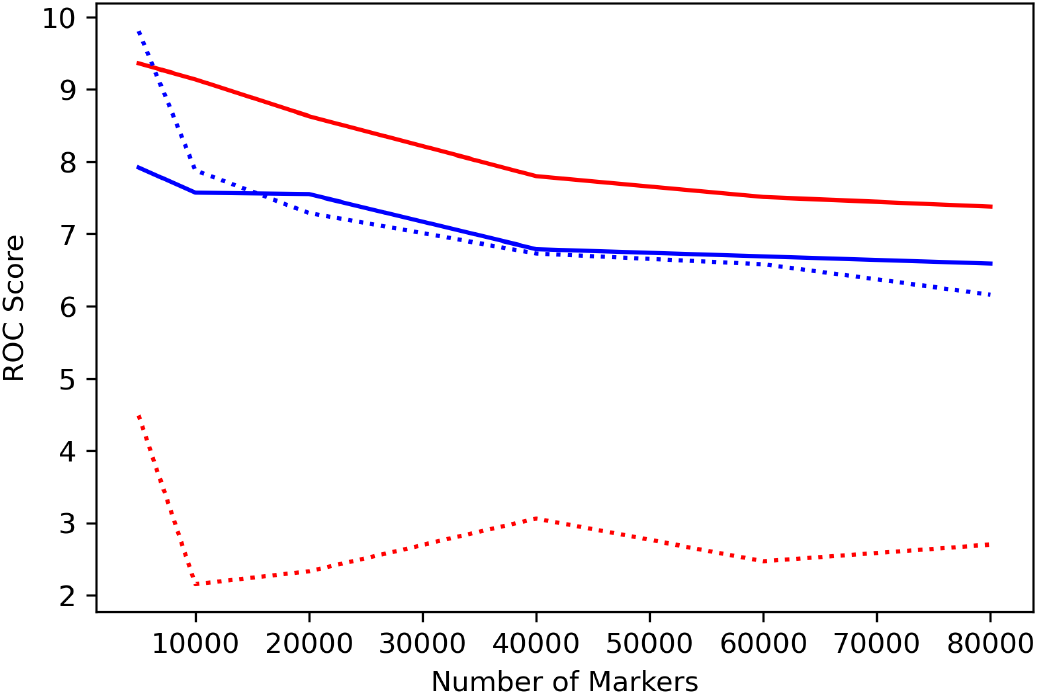
The *ROC score* of a GWAS experiment under varying number of markers, with blue lines representing Bonferroni correction and red lines representing BH-FDR. Solid lines representing independent markers whereas dashed lines representing correlated markers. The sample size is maintained at 2000, the number of QTL is maintained at 100, and the average QTL effect size (*γ*) and allele frequencies distribution shape parameter (*β*) maintained at 0.4 and 0.5 respectively.

Besides the number of markers used, sample size also has a significant effect on the *ROC score* for both multiple testing correction methods (Figure 15). When the markers are independent, compared to Bonferroni correction, BH-FDR has somewhat higher *ROC score* in all sample sizes tested, although this observation is more notable for very small sample size (N=200) or for the larger sample size (N=5000). The presence of correlation changes the trend however; for N=200, the *ROC score* for BH-FDR is higher than for Bonferroni, but with sample size of 800 and larger the *ROC score* of Bonferroni is higher than that of BH-FDR, and with sample size larger than 1400, the *ROC score* actually decreases with larger sample size for BH-FDR. This is attributable to an increased number of of false positives for BH-FDR with large sample sizes.

**Figure 15.**
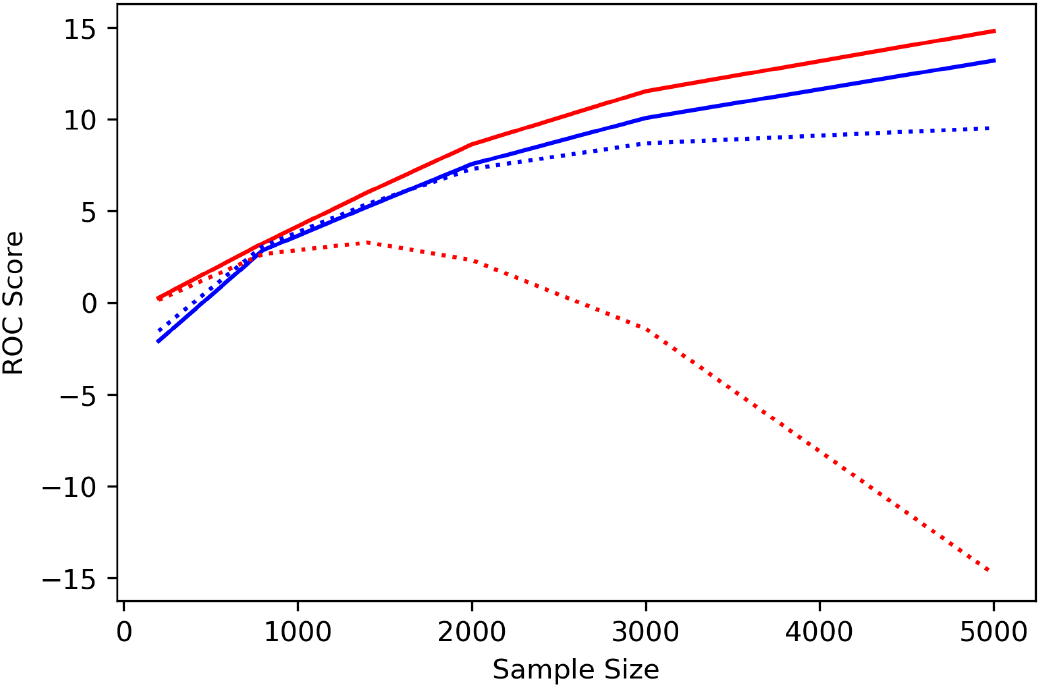
*ROC score* of a GWAS experiment under varying sample sizes used in GWAS for Bonferroni correction (blue lines) and BH-FDR (red lines) for independent markers (solid lines) and correlated markers (dashed lines). The number of markers and QTL is set at 20k and 100 respectively, and the average QTL effect size (*γ*) and allele frequencies distribution shape parameter (*β*) maintained at 0.4 and 0.5 respectively.

For independent markers, the *ROC score* for both BH-FDR and Bonferroni correction would initially increase, but when the number of QTL exceeds 100 there is a slow decline in the *ROC score*. This can be attributed to a decline in power with increasingly large number of QTL that have smaller effect. For correlated markers, the *ROC score* of Bonferroni correction followed a similar trend as with independent markers, but the trend is different for BH-FDR. When the number of QTLs is small (less than 200) the *ROC score* went below that of Bonferroni correction, but with further increase in number of QTL, the *ROC score* with BH-FDR became significantly higher than that of Bonferroni (Figure 16).

**Figure 16.**
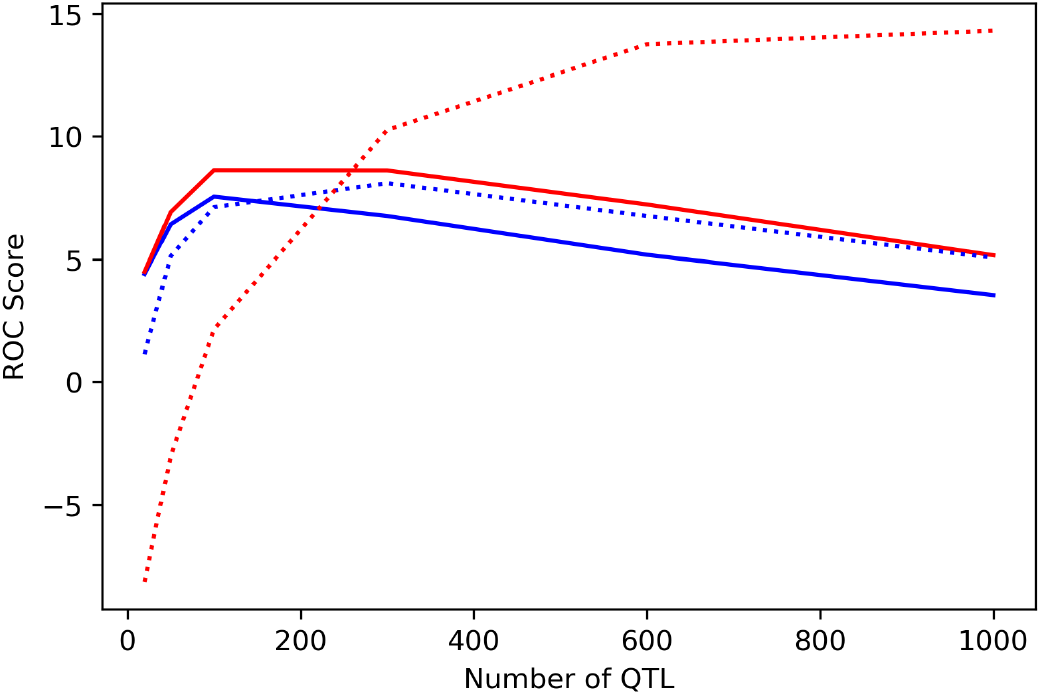
*ROC score* of a GWAS experiment under varying number of QTLs, with blue lines representing Bonferroni correction and red lines representing BH-FDR. Solid lines represent independent markers and dashed lines represent correlated markers. The sample size is maintained at 2000, the number of markers at 20k. The average QTL effect size (*γ*) and allele frequencies distribution shape parameter (*β*) maintained at 0.4 and 0.5 respectively.

With increased numbers of QTL with large effect sizes (i.e. a high average QTL effect sizes), the *ROC score* for both multiple testing correction method increases (Figure 17). This can be attributed to an increase in power with increasingly large average QTL effect sizes. When the markers are independent, BH-FDR has again a higher *ROC score* than Bonferroni for all parameter values, although the increase is more significant with a larger average QTL effect size. This trend flipped when the markers are correlated however; while BH-FDR has high power in detecting QTLs, the massive increase in number of false positives decreased the *ROC score* to that below of Bonferroni correction. Correlation has a less significant effect on the *ROC score* for Bonferroni correction.

**Figure 17.**
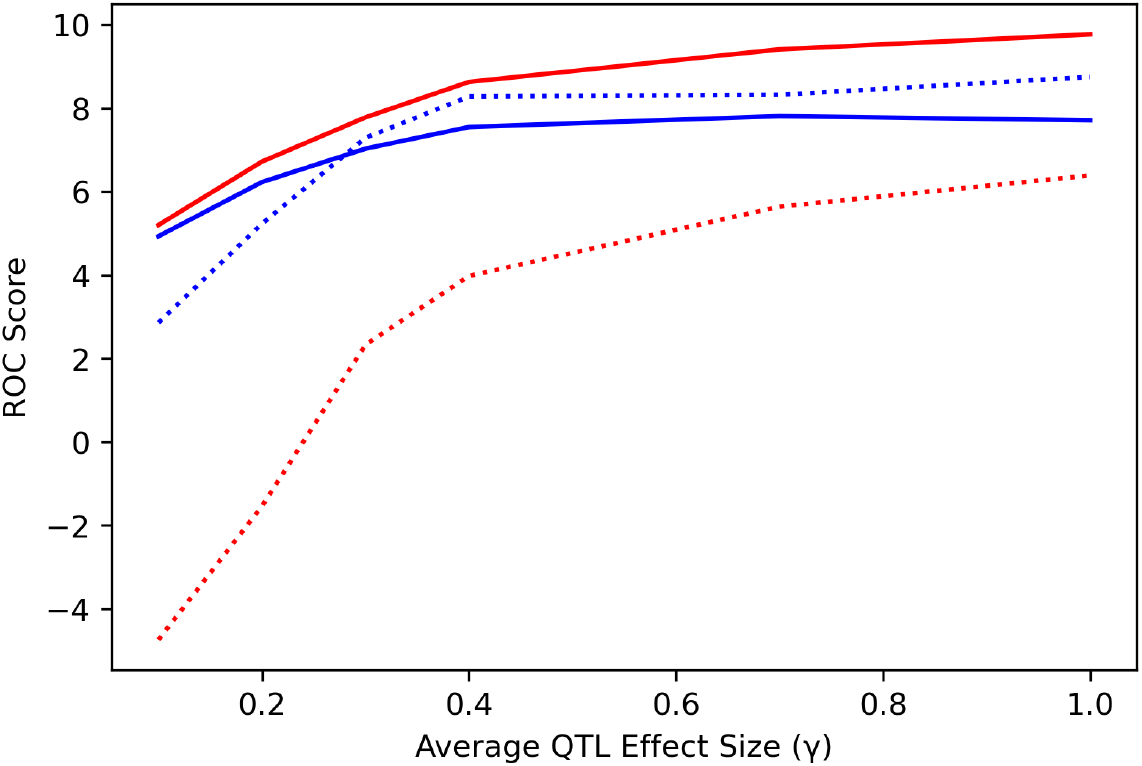
*ROC score* of a GWAS under varying shape parameters for allele substitution effects for Bonferroni correction (blue lines) and BH-FDR (red lines) and for independent markers (solid lines) and correlated markers (dashed lines). The sample size is maintained at 2000, the number of markers and QTL at 20k and 100 respectively, and the shape parameter for allele frequency distribution (*β*) set at 0.4.

## Discussion

In this experiment the effects of parameters on the threshold of Bonferroni correction and BH-FDR, as well as its associated power and false positive rate, were tested. Unlike BH-FDR, which has its threshold affected by various parameters, none of these parameters have an effect on the threshold of the Bonferroni correction, with the exception of number of SNP markers. This is due to how the threshold is calculated; with the number of SNP markers being the only variable for the calculation of threshold for Bonferroni correction (equation (5)). The threshold of BH-FDR also depends on the distribution of the ranked p-values of the markers (i.e. rank “k” in equation (6)). Due to this, any parameters that could affect the distribution of p-values would have an effect on the threshold from BH-FDR. Parameter values that would increase the -log(p-value) of the markers would increase the value of point k and thus decrease the stepwise threshold from (6), thus decreasing the stringency of the threshold. Conversely parameters that decrease that -log(p-value) would decrease the value of point k and thus increase the threshold stringency. For example, increasing the sample size of the GWAS increases the test statistic of the marker and thus increases the value of the -log(p-value). This causes an increase in the value of k and thus decreases the stringency of threshold. Conversely a trait with a large number of QTL would decrease the proportion of phenotypic variance explained of any given QTL, and this would increase the p-values and thus the stringency of the threshold of BH-FDR. Correlations between markers would also causes the “bleeding” of effect sizes from the true markers into the neighbouring null markers, and this would produce a peak of true QTL with several neighbouring null markers flanking the peak (Figure 18). This increases the p-values of the neighbouring null markers and thus decreases the threshold of BH-FDR.

**Figure 18.**
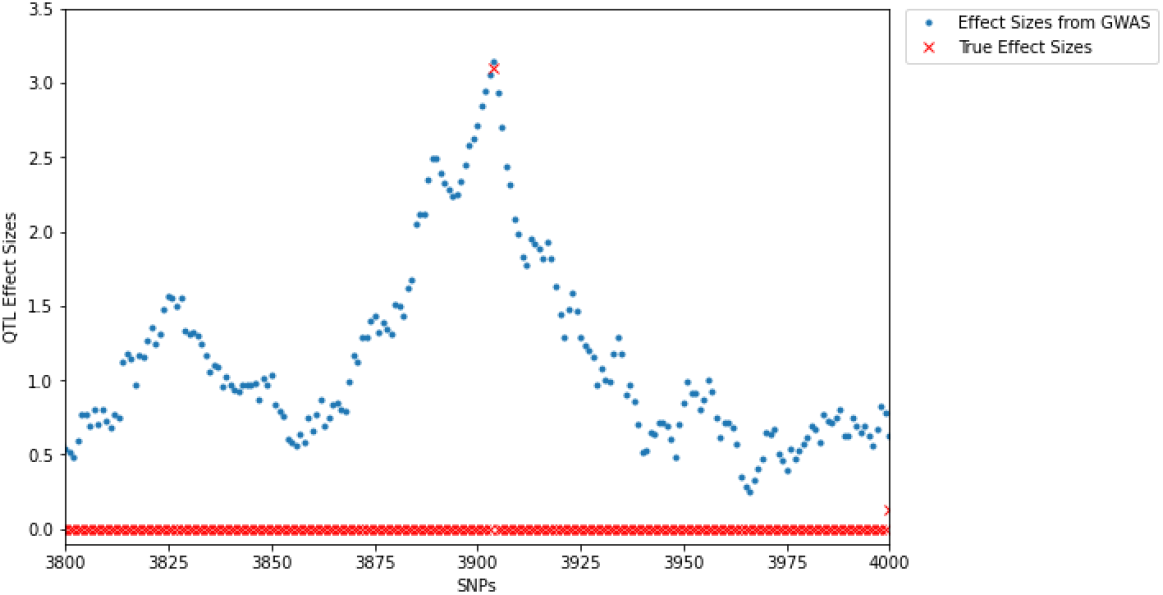
The estimated marker effect size of a peak in a correlated genotypes, showing the effect of correlation on the null markers that flanked the true QTL marker. The marker pair correlation is set at *r*^2^ = 0.95.

The effects of these changes in the threshold of both correction methods would affect the power and false positive rate of a GWAS, with a decreased stringency in threshold increases its power and false positive rate and vice versa. For example, increasing the number of markers caused the threshold to become more stringent as it needs to exclude the additional null markers. This increased stringency however also has the effect of decreasing the number of true positives and thus the power. As the threshold would increase in a logarithmic fashion with an increase in the number of markers, the power would also decreases in a similar fashion, approaching zero as none of the true QTL had its p-value exceed the extremely stringent threshold. This could be an issue for Whole Genome Sequencing (WGS) data, where Tam *et al*. (2019) warned the exacerbation of decline in power due to the overconservative threshold, especially when Bonferroni correction is used. In this situation BH-FDR might serve as a better alternative. A larger number of markers tested also does not necessarily increase the power, as it might increase the number of null markers utilized in a GWAS experiment, and increase the required stringency of a threshold.. In fact, we saw a decrease in power with a larger number of markers tested in both correlated and uncorrelated markers. Conversely increasing the sample size would increase the -log(p-value) of the true markers, making them more likely to be detected. This increases the number of true positives logarithmically, thus increasing the power of GWAS. This suggests that increasing the sample size could be more important than increasing the number of markers used in a GWAS experiment.

On the other end of the spectrum, the use of small sample size significantly increased the number of false positives and decreases the number of true positives. This is due to the fact that observations made from a small sample size can often be explained by a larger number of predictors (i.e. SNPs markers), which causes the null markers that have its combination of genotypic values coincided with those of true markers to have an elevated p-value, contributing to the false positive rate (Forstmeier *et al*. 2017). Combined with the reduced number of true positives, this would mean a GWAS with small sample size would have low power and high false positive rate. This observation is also supported by the low *ROC score* for small sample sizes, and this elevated the number of false positives, especially with increasingly large number of SNPs. As expected, the result of this experiment have casted doubt on the validity of the results obtained from studies with a small sample size, especially for those with high marker density.

Besides the experimental design parameters, the distribution of the QTL effect size of the trait studied would also affect the threshold, power and false positive rate of a multiple testing correction method in a GWAS experiment. This agrees with the study of Panagiotou and Ioannidis (2012) which stated that the most suitable threshold used in a GWAS experiment might vary for different populations and genetic architecture of the trait. This could mean that a threshold from one study might not be suitable for another association study. Indeed, this can be observed with the decreased number of true positive as well as increased number of false positives for a trait with a small number of QTL with large effect sizes (small average QTL effect size in this study). While in theory this observation could be used to calculate the threshold optimized for the trait in study, in practice this might not be possible as it requires information on the underlying QTL effect size distribution. Previous works such as those published by Park *et al*. (2011), Hall *et al*. (2016) and Zhang *et al*. (2018) have attempted to estimate such distribution using various approaches, although these algorithms assumed the QTL effect sizes followed exponential or normal distribution, and the effect of violation of such assumption (as an example, the QTL effect size followed a gamma distribution) remained untested. Future work should be thereby focus on an algorithm for robust estimation of QTL effect size distribution, and its incorporation into the calculation of optimal threshold for a GWAS experiment.

Panagiotou and Ioannidis (2012) also commented that correlated markers constituted a major source of uncertainty in the suitability of a threshold used in a GWAS experiment. Our results show that this is a valid concern. Due to the “bleeding” effect of the effect sizes, correlation between markers significantly increased the number of false positive in a GWAS experiment. This would be a valid concern with increasingly large number of markers used. As markers become increasingly dense, they become less well separated to one another and they no longer inherited independently, producing linkage disequilibrium between markers (Cheverud 2001; Falconer 1989). It is expected that maximal linkage disequilibrium would be observed for WGS data, which made high degree of correlation between markers unavoidable (Pengelly *et al*. 2015).

While correlations between markers led to an increased number of false positives for BH-FDR, and increase in number of QTL also led to a decline in the power of GWAS, such that increase in number of QTLs had led to an increase in the absolute number of true positives and a decrease in number of false positives. This led to an increase in the *ROC score* and a decrease in the false positive rate. This could be attributed to an increase in threshold for BH-FDR with large number of QTLs associated with a trait, as well as the reduced proportion of variance explained by each of the QTL. Such a decrease in false positive rate and increase in *ROC score* was neither observed for Bonferroni correction, nor for independent markers. While this initially suggested that BH-FDR might be more suitable correction method compare to those of Bonferroni for a polygenic trait, especially with correlated marker, it does come with a caveat: the actual false positive rate in BH-FDR is significantly higher than that of Bonferroni; the false positive rate from FPU is 0.095 under default conditions for BH-FDR, compared to 0.006 for Bonferroni. Thus the actual suitability of the correction methods depends on the priority of the experiment; if the priority of the GWAS is to be placed on the explanatory power of the GWAS, then BH-FDR might serve as a better choice, but if the priority is to increase the specificity (i.e. number of false positives out of all null markers) of the GWAS, then Bonferroni might be preferred.

Rather than choosing one multiple testing correction methods over the other, perhaps a better alternative is to modify the methods so that they could take into account correlation between markers. One such method is a Bonferroni correction that utilized “effective number of independent markers” instead of raw number of SNP markers, similar to those suggested by Cheverud (2001) and Nyholt (2004). This might serve as a promising route for an increased power. The calculation of effective number of independent markers utilized the variance of eigenvalues obtained from the marker correlation matrix to adjust the “Number of SNPs” in equation (3), thus yielding a less stringent threshold. One downside of this method however is that the calculation of a very large marker correlation matrix is memory and computationally demanding, which might not be feasible with large dataset. Modification of the original methods might thus be required. Studies on the effect of such adjustment of “Number of SNPs” on the false positive rate of GWAS is also lacking as well. Attempts to modify the BH-FDR so that it could take into account correlation between markers had also been done by Benjamini and Yekutieli (2001), and this might serve as a better alternative than BH-FDR.

While correlation between markers is expected to be at its strongest with WGS data, the effects of correlation on setting the threshold in a multiple testing correction method cannot be ignored in GWAS experiment where WGS data is not used. With pairwise marker correlation as high as 0.8, it has caused such a significant decline in the stringency of the threshold for BH-FDR that it fails to control the number of false positive and thus the false discovery rate at 0.05 in all parameter value tested. This highlighted the importance of selecting the appropriate multiple testing correction method in a GWAS experiment. The results from this experiment has also ran contrary to the claims from Benjamini and Yekutieli (2001) on the validity of BH-FDR in correlated marker array. This experiment highlighted the unsuitability of BH-FDR with high density marker arrays, which is to be expected in a real GWAS experiment, especially when the markers are not sufficiently dense. In this situation Bonferroni correction was shown to be more capable of maintaining the number of false positives.

An important note for this experiment, which would serve as a caveat, is how the number of FPU and FPC is determined. It should be noted that the cut-off point between FPU and FPC (i.e. *r*^2^ = 0.1) is arbitrary, and changing said cut-off point would affect the number of FPU and FPC. The rationale of using this cut-off point is to differentiate the “false positives” that is caused by correlation with true QTL from those that actually caused by the varying parameter values. This distinction is important in the context of increasing the sample size of the GWAS. While the naïve result of this experiment suggested that increasing the sample size would increase the false positive rate of a GWAS, this is more likely the effect of choosing a certain cut-off point for FPC and FPU, as increasing the sample size would not only decreases the required effect size detectable by the GWAS, but also the required correlation between the marker and the QTL. Given a probability of detection of a QTL, the required correlation between a marker and QTL to achieve said probability of detection decreases with increasingly large sample size. This is best illustrated by Spencer *et al*. (2009) who had provided the following proportionality between the test statistics for the detection of a QTL and correlation between QTL and marker:

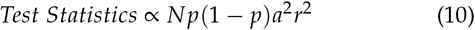

Where *N* is the sample size, *p* is the allele frequency, *a* being the QTL effect size and *r*^2^ being the correlation between QTL and marker. This proportionality suggests that for a given a test statistic value, as *N* approaches infinity, the *r*^2^ required for the test statistic to reach said value would approaches zero. Thus, as long as a marker has a nonzero correlation with any of the QTL, regardless how small the correlation is, there would be a finite sample size required. Taking this to extreme, this could cause a GWAS to declare excessively large number of null markers to be positive, even if those markers are minimally correlated to the QTL. This could be the situation observed by Jiang *et al*. (2019), who have utilized 294,079 animals in their GWAS experiment, and with that sample size the experiment declared 27% of all markers as positives (15215/57067 = 0.267). In the context of multiple testing correction methods, this would also suggest that a GWAS with large sample size could afford a more stringent threshold, such as those suggested by Bonferroni correction.

Given that differing genetic architecture of a trait and experimental designs would affect the suitability of the threshold of a multiple testing method, an algorithm that could test the suitability of such threshold would be desirable. One possible method of testing the appropriateness of the multiple testing correction method is to use the *ROC score*. As defined by equations (7), (8) and (9), a low *ROC score* could either be caused by low power, which would be associated with an overconservative threshold, or with high false positive rate, which would be associated with overly lenient threshold. Only a threshold that could provide a good balance between true and false positives that would have a high *ROC score*. Results from this experiment suggested that when the markers are independent, BH-FDR provided a better balance between power and false positive rate for all parameter values tested when markers are not correlated, but for correlated markers, Bonferroni correction consistently provided a better balance between power and false positive rate for all parameter value tested except for highly polygenic trait (i.e. trait with large number of QTL).

Given the relationship between the *ROC score* and the optimality of the threshold, another potential route for further study is to establish an algorithm that could find an optimal threshold that would maximize the *ROC score* based on certain parameters related to experimental design and genetic architecture of the trait in study, similar to what is suggested by Habibzadeh *et al*. (2016) in finding a threshold that balances the power and false positive rate in a clinical test, and de Smet *et al*. (2004) had used such an algorithm in balancing true and false positives in a gene expression experiment. Along with suitable modification, such as taking into account the effect of correlation between markers, a similar algorithm could be suggested to be used in a GWAS experiment. A major obstacle for this route is the requirement of prior information on the underlying QTL effect size distribution, which further emphasize the importance of a robust algorithm to estimate it.

In conclusion this experiment suggested that power and false positive rate in a GWAS experiment is affected by the choice of the multiple testing correction method, the experimental parameters such as sample size and number of markers, and the genetic architecture parameters of the trait studied. For independent markers, BH-FDR provided a better balance between the true and false positives for all parameter values, but for correlated markers, Bonferroni correction did provide a better balance between true and false positives. The only exception where BH-FDR provided a better balance between true and false positive with correlated markers is when the trait is highly polygenic, and even so with the caveat of increased false positive rate. This experiment had also suggested that increasing the number of markers used in an experiment would not necessarily increase the power of GWAS, but increasing the sample size would increase the power and decrease the false positive rate of GWAS. Our study also showed the importance of having large sample size if large number of markers is to be used in a GWAS experiment, which would be crucial if WGS data is to be used in a GWAS experiment, as a genotype array of such high density would inevitably and excessively increase the stringency of the threshold, necessitating a larger sample size. Future work should focus on a robust algorithm to estimate the QTL effect size distribution, and using it to calculate an optimal threshold that could balance the power and false positive rate under arbitrary experimental designs and genetic model of the trait studied in a GWAS experiment.

## Conflict of Interest

We declare no conflict of interest for this publication.

